# Keystone Bacterial Taxa Drive Denitrification and N_2_O Emission via Adaptive Genomic and Metabolic Strategies in Contrasting Agricultural Soils

**DOI:** 10.1101/2025.07.17.665313

**Authors:** Siyu Yu, Qiaoyu Wu, Yimin Ma, Xiaojun Zhang

**Affiliations:** State Key Laboratory of Microbial Metabolism, Joint International Research Laboratory of Metabolic & Developmental Sciences, and School of Life Sciences & Biotechnology, Shanghai Jiao Tong University, Shanghai, 200240, China; State Key Laboratory of Pollution Control and Resources Reuse, Shanghai Institute of Pollution Control and Ecological Security, College of Environmental Science and Engineering, Tongji University, Shanghai 200092, China

**Keywords:** Keystone taxa, Microbial co-occurrence network, Nitrous oxide (N_2_O) emission, Denitrification, Soil

## Abstract

Soil denitrification mediated by microbial communities is a major driver of nitrous oxide (N₂O) emissions, yet the regulatory roles of keystone taxa remain largely unexplored, particularly under distinct edaphic conditions. Black soil (BS) and fluvo-aquic soil (FS), two representative agricultural soils in China, exhibit contrasting N₂O emission potentials, providing a unique model system for disentangling microbial mechanisms underlying soil-specific denitrification dynamics. Here, we integrated microbial co-occurrence networks, metagenomics, and functional phenotyping to identify and characterize keystone bacterial taxa involved in denitrification across the above mentioned two contrasting agricultural soils, BS and FS. Structural equation modeling (SEM) and correlation analyses revealed that keystone taxa are significantly associated with soil denitrification rates and N₂O emission patterns. Among these, *Ramlibacter* ASV104 was identified as a conserved keystone in both soils, exhibiting genomic potential for key denitrification genes (*nirK*/*nirS*/*norB*/*nosZ*) to mediate N_2_O turnover. In contrast, *Ensifer* ASV205 was a FS-specific keystone taxon, exhibiting strain-level niche specialization. Comparative genomics revealed that variations in denitrification gene composition and carbon-nitrogen metabolism enable different *Ensifer* strains to function either as N₂O sources or sinks. This functional divergence was linked to their genomic plasticity and growth strategies under nutrient-rich conditions. Our findings demonstrate that soil-specific denitrification processes and N₂O emissions are governed by keystone taxa through adaptive genomic and metabolic strategies shaped by environmental filtering. This study improves our understanding of the microbial mechanisms driving N₂O emissions and provides a foundation for future strategies targeting microbial taxa to mitigate N₂O emissions in agricultural soils.

**IMPORTANCE:** This study provides new insights into the ecological complexity underlying soil N₂O emissions by integrating strain-level diversity, genomic attributes, and microbial interaction networks. Key microbial taxa involved in denitrification and N₂O emissions were identified in black soil and fluvo-aquic soil. By combining network-based ecological modeling and functional assays, it is revealed that taxa such as *Ramlibacter* ASV104 and *Ensifer* ASV205 play significant roles in regulating N₂O emissions. Strain-level functional diversity within the keystone taxa of *Ensifer* ASV205 highlights their varying contributions to nitrogen cycling. Our findings provide a deeper understanding of the microbial mechanisms governing N₂O emissions, offering valuable insights for developing microbial-based strategies to mitigate N₂O emissions in agricultural soils in the future.

## INTRODUCTION

Soil microorganisms are fundamental drivers of terrestrial ecosystem functioning, playing vital roles in regulating carbon and nitrogen cycling, greenhouse gas emissions, and nutrient transformation, thereby directly or indirectly influencing soil productivity and ecological balance (1). These microbial communities are taxonomically diverse and highly complex, maintaining soil stability and multifunctionality through dynamic interaction networks (2). Within these intricate networks, keystone taxa act as ecological “hubs” due to their high connectivity. They can profoundly shape community structure and ecosystem functions via metabolic interactions, resource competition, and signal transduction (3–7). The removal of such taxa can lead to substantial shifts in microbial community composition and functionality, often led to malfunction or even collapse of specific ecosystem (4). Notably, previous studies have demonstrated that keystone taxa explain microbiome compositional turnover more effectively than the combined contributions of all taxa (8), and their presence enhances soil ecosystem multifunctionality (2). The regulation of community structure and function by keystone species is influenced by environmental heterogeneity and changes in nutrient resources (9, 10). Recent analyses of soil microbial network structures and keystone species dynamics have further elucidated the microbial mechanisms through which environmental factors and agricultural management influence soil functions (11–13). However, the identification and characterization of keystone taxa within such complex microbial communities remain a significant challenge due to lack of experimental verification from microbial culture. Fine scale diversity is another obstacle to describe the keystone taxa in complicated soil environment. More and more studies have shown that different strains of the same species exhibit varying ecological functions under different environmental conditions (14, 15).

Denitrification is a critical microbial process for soil nitrogen cycling that facilitates the stepwise reduction of nitrate (NO₃⁻) and nitrite (NO₂⁻) through nitric oxide (NO) and nitrous oxide (N₂O) to dinitrogen (N₂). The complete denitrification pathway is catalyzed by enzymes encoded by *narG*/*napA*, *nirS*/*nirK*, *norB*, and *nosZ*, respectively(16). Nitrous oxide (N₂O) is a long-lived trace gas that contributes significantly to global warming and stratospheric ozone destruction. It is recognized as one of the most critical greenhouse gases, second only to carbon dioxide (CO₂) and methane (CH₄)(17). Since the pre-industrial era, atmospheric N_2_O concentrations have increased from 270 parts per billion (ppb) in 1750 to 331 ppb in 2018 (18). Agriculture is the largest source of anthropogenic N₂O emissions (19), with cropland soils estimated to contribute over 80% of global anthropogenic N₂O emissions (20). The increasing global population and rising dependence on agricultural fertilizers have collectively led to a worrying increase in anthropogenic N₂O emissions, which will further aggravate climate change and ozone layer depletion(21). Notably, only the nitrous oxide reductase (NosZ) in the denitrification process catalyzes the reduction of N₂O to N₂. As a result, denitrifying microorganisms containing *nosZ* are the sole potential sinks for N₂O (22). And N₂O net emissions in soils depend on the overall denitrification rate and N₂O/N₂ product ratio (23).

It has been suggested that environmental factors, human activities, and functional microbial communities can explain approximately 59% to 68% of N₂O emissions (24). In addition, N₂O emissions and their response to environmental changes are also influenced by the interactions both within and between microbial groups in the soil microbiome (25). The population abundance, community structure, and functional activity of microorganisms involved in denitrification processes are critical factors in regulating N₂O emissions (26). Microbial network analysis can provide important and clear insights into microbial communities, particularly in explaining microbial interactions and identifying the functional key microbial taxa (27, 28). Focusing on keystone taxa is expected to more effectively decipher the mechanisms through which microbial communities modulate soil denitrification and N₂O emissions. This approach also enables the identification of the pivotal roles played by individual species within the microbiome, offering deeper insights into their functional contributions to nitrogen cycling processes.(6). To date, relatively few studies have utilized network analysis methods to infer potential keystone taxa closely associated with denitrification processes and N₂O emissions across diverse ecosystems(7, 29, 30). However, these only existed studies have neglected the functional diversity among different strains of keystone species. Given the high phenotypic diversity of denitrifiers (31), these studies potentially led to incomplete interpretations of the functional regulatory mechanisms governing soil microbiomes.

Black soil and fluvo-aquic soil are widely distributed and highly representative soil types in China, playing critical roles in agricultural production. Black soil, characterized by high fertility and abundant organic matter, is predominantly distributed in northeastern regions of China. In contrast, fluvo-aquic soil is main soil type in the North China Plain, where periodic flooding and spatiotemporal variability in nutrient supply shape their ecological dynamics. Our previous studies have revealed that the distinct physicochemical properties of black soil and fluvo-aquic soil have shaped adaptive microbial communities, thereby modulating denitrification processes and N₂O flux dynamics (32, 33). However, the mechanisms how keystone taxa drive these soil-specific functional phenotype remain poorly understood. This study aims to identify highly connected, functionally keystone taxa associated with soil denitrification and N₂O emissions through a systematic investigation of microbial network structures and denitrifying communities in these two distinct soil types. We constructed microbial co-occurrence networks to identify keystone taxa. And further explored the denitrification and N₂O-reducing capabilities of these taxa after employing metagenome-assembled genome (MAG) analysis, and extensive cultivation of culturable bacterial taxa. Accurately identifying and functionally characterizing these key microorganisms will enhance our understanding of nitrogen cycling in soils and support the development of sustainable agricultural practices.

## RESULTS

### Denitrifying communities and associated genes in two type soils

The species composition and functional gene abundance related to nitrogen transformation pathways in black soil and fluvo-aquic soil were compared using metagenomic data. The structure of dominant denitrifying species in black soil and fluvo-aquic soil was similar, with 12 major denitrifying genera including 12-FULL-67-14b, UBA5189, *Luteimonas B*, *Ramlibacter*, QHWT01, *Spingomicrobium*, Gp6-AA40, PSRF01, etc. Many denitrifying species remain unclassified at the genus level (Fig. 1a). A total of 30,094 microbial species were shared by both soils, primarily belonging to the phyla *Pseudomonadota* (32.9%), *Actinomycetota* (14.7%), *Bacteroidota* (9.1%), and *Chloroflexota* (4.6%). Black soil contains 3,136 unique species, while fluvo-aquic soil has 2,864 unique species. Specifically, 1,041 denitrifying species were shared by both soils. Black soil contains 112 unique denitrifying species, while fluvo-aquic soil has 107 unique species (Fig. 1b). It was found that fluvo-aquic soil exhibited higher abundance of functional genes associated with organic carbon degradation and synthesis, anaerobic ammonium oxidation, assimilatory nitrate reduction, nitrification, denitrification, and dissimilatory nitrate reduction to ammonium (DNRA) compared to black soil (Fig. 1c). The abundances of both N₂O-producing functional genes (*nirK*/*norB*) and the N₂O-reducing gene (*nosZ*) in fluvo-aquic soil were higher than those in black soil (Fig. 1d).

**Fig. 1.**
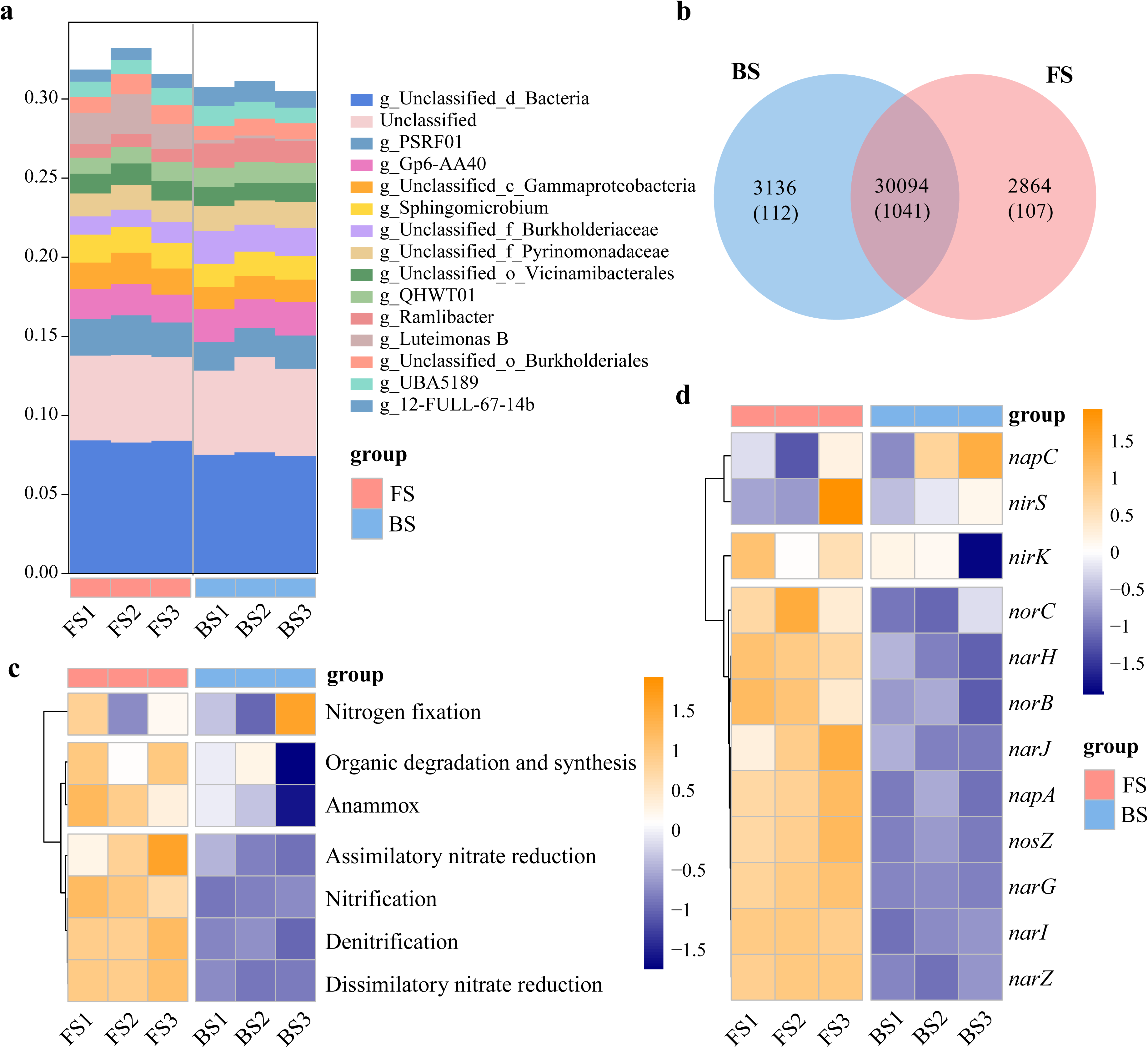
Denitrifying communities and abundance of nitrogen transformation functional genes in black soil (BS) and fluvo-aquic soil (FS) based on metagenomic data. (a) Relative abundance of dominant genera related to denitrification. (b) Venn diagram showing the shared and unique species between the two soils. Numbers in parentheses represent the number of shared and unique denitrifying bacterial species between the two soils. (c) Heatmap of the abundance (TPM) of different nitrogen transformation pathways. (d) Heatmap of abundance (TPM) of denitrification functional genes.

### Diversity of denitrifying bacteria in two type of soils

The specific denitrification function and N₂O reduction potential of network hub species was investigated by combining strain isolation and metagenomic binning. Metagenomic data from BS and FS samples were co-assembled and binned, resulting in a total of 261 metagenome-assembled genomes (MAGs). A total of 63 MAGs were retained following redundancy removal, meeting the criteria of completeness >50% and contamination <10%. According to the phylogenetic analysis based on the Genome Taxonomy Database (GTDB), 63 bacterial MAGs belonging to 9 different phyla were obtained. Of these, 31.7% were affiliated with *Pseudomonadota*, followed by *Acidobacteriota* (30.2%) and *Gemmatimonadota* (19.0%) (Fig. 2).

**Fig. 2.**
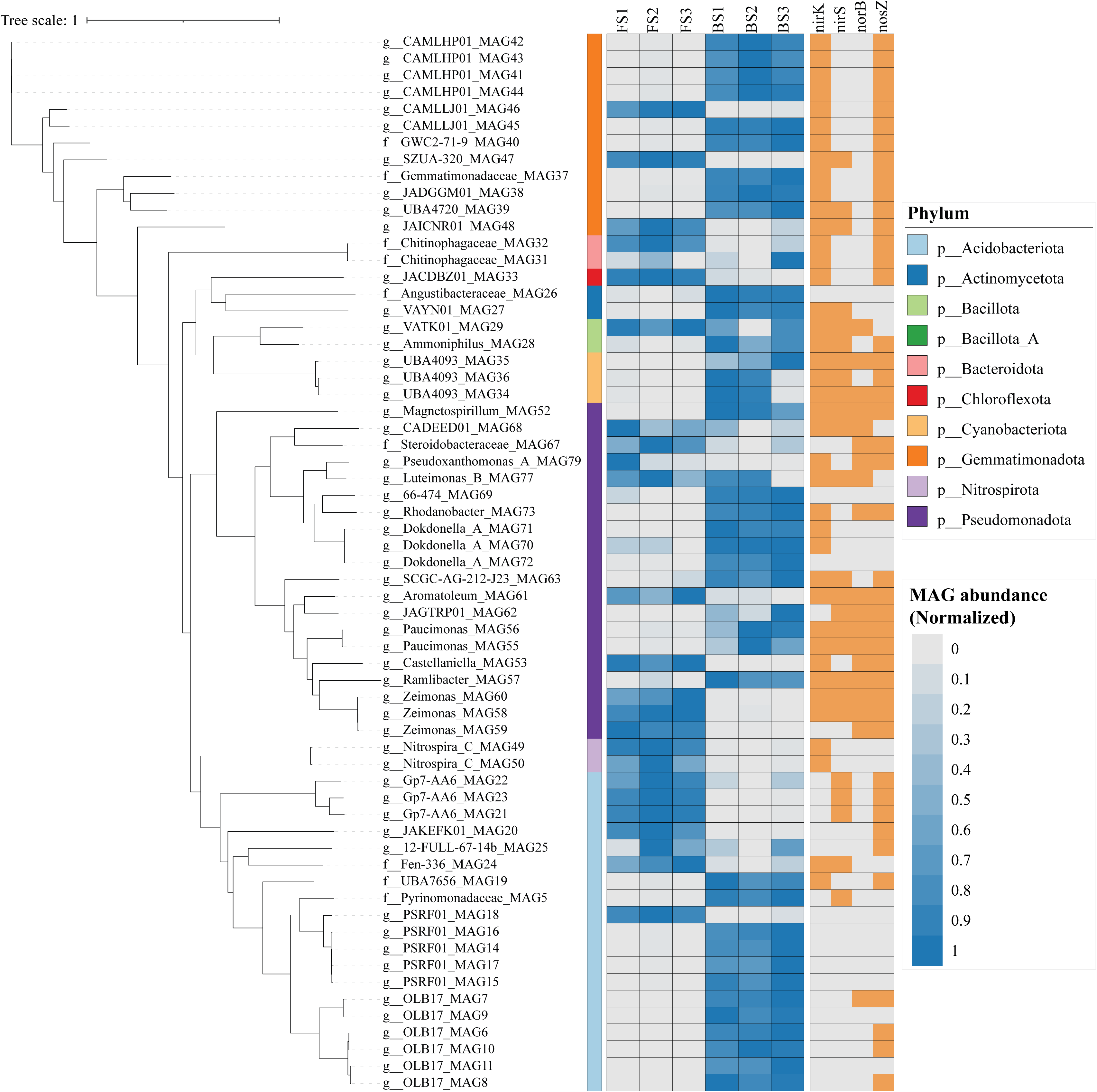
The phylogeny, nitrogen cycling genes and abundance of MAGs recovered from metagenomic data of BS and FS. These MAGs were selected using CheckM (completeness >50%, contamination <10%) and identified using the Genome Taxonomy Database. The most left panel is the phylogenetic tree of MAGs. The heatmap on the middle panel illustrates the relative abundance of Metagenome-Assembled Genomes (MAGs) in BS and FS. The orange squares on the right indicates the presence or absence of denitrification genes (*nirK*, *nirS*, *norB*, *nosZ*) within these MAGs.

Denitrifying MAGs in black soil predominantly belonged to CAMLHP01, CAMLLJ01, *Rhodanobacter*, *Castellaniella*, *Ramlibacter*, *Zeimonas*, *Aromatoleum*, and *Nitrospiro*. The TPM-normalized abundance values highlight clear differences in MAG enrichment between the two soil types. Several MAGs, including those affiliated with *Castellaniella* (MAG53), *Aromatoleum* (MAG61), *Luteimonas_B* (MAG77), and *Zeimonas* (MAG40), exhibited higher abundance in FS, while *Ramlibacter* (MAG57) and *Rhodanobacter* (MAG55) were more dominant in BS. These patterns suggest that specific denitrifying populations are differentially selected by the environmental conditions of each soil type, contributing to the observed variation in nitrogen cycling potential. MAGs harboring *nosZ* but lacking *norB* gene were affiliated with CAMLHP01, CAMLLJ01, GWC2-71-9, SZUA-320, JADGGM01, UBS4720, JAICAR01, JP7-AA6, and OLB17. They might exist as N_2_O sink in two soils. Differential abundance patterns of CAMLLJ01-affiliated MAGs were observed between black soil and fluvo-aquic soil (Fig. 2). These taxonomically closely related strains may occupy distinct ecological niches and exhibit divergent functional roles in the two soils.

By implementing both aerobic and anaerobic cultivation protocols, a total of 719 bacterial strains were successfully isolated from black soil and fluvo-aquic soil. To investigate the taxonomic composition and functional potential of cultivable denitrifiers, a total of 135 unique ASVs were obtained from high-throughput sequencing of bacterial isolates from BS and FS. Each ASV represented a unique 16S rRNA variant, corresponding to distinct bacterial strains. These isolates were distributed across diverse taxonomic clades, including *Bacillus*, *Arthrobacter*, *Ensifer*, *Mesorhizobium*, and *Paenibacillus* (Fig. 3a). 22 bacterial genera were isolated from BS and 33 genera were isolated from FS. PCR-based screening targeting denitrification functional genes (*nirK*, *nirS*, *norB*, *nosZ* Ⅰ*, nosZ* Ⅱ) identified potential denitrifying strains belonging to *Agromyces*, *Arthrobacter*, *Azospirillum*, *Bacillus*, *Bosea*, *Bradyrhizobium*, *Castellaniella*, *Cellulomonas*, *Cupriavidus*, *Cellulosimicrobium*, *Diaphorobacter*, *Ensifer*, *Lysobacter*, *Mesorhizobium*, *Microbacterium*, *Microvirga*, *Nocardia*, *Paenibacillus*, *Pseudomonas*, *Phycicoccus*, *Rhizobium*, *Streptomyces*, and *Sphingopyxis*. Strains with N₂O reduction potential predominantly belonged to the phylum *Pseudomonadota* and were mainly affiliated with the genera *Azospirillum*, *Bacillus*, *Ensifer*, *Mesorhizobium*, *Microvirga*, *Castellaniella*, *Cupriavidus*, *Paenibacillus*, *Phyllobacterium*, *Pseudaminobacter*, *Streptomyces* and *Sphingopyxis.* (Fig. 3a). To further evaluate genus-level functional trends of isolates, ASVs from the top 9 most diverse genera were classified as non-denitrifiers, denitrifiers lacking *nosZ* and N₂O reducers based on functional gene screening. *Bacillus* and *Arthrobacter* exhibited the highest strain-level diversity, each represented over 15 unique ASVs (Fig. 3b).

**Fig. 3.**
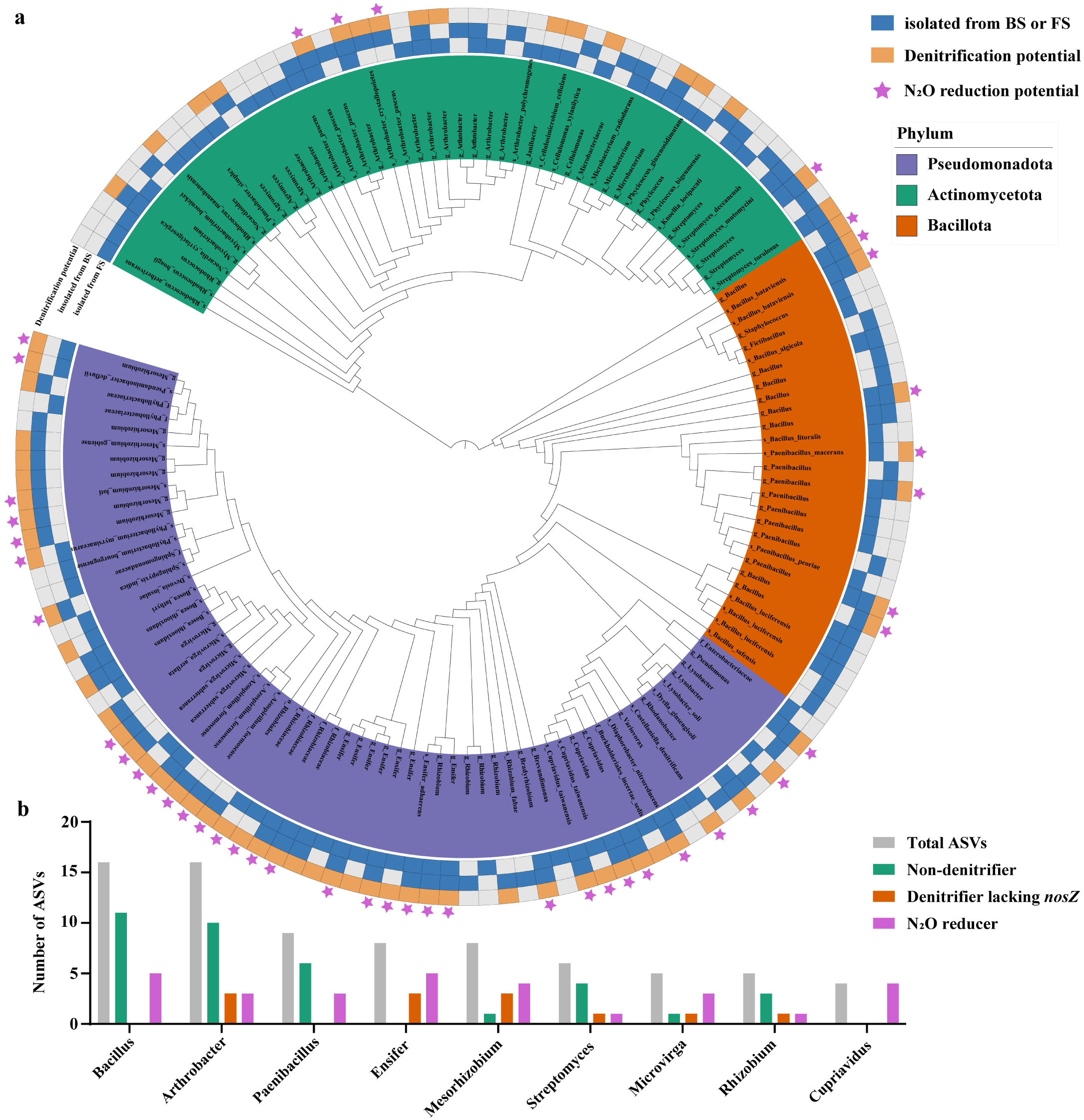
Phylogentic tree, taxonomic distribution and function of cultivated bacteria. **(a)** Maximum-likelihood phylogenetic tree of de-replicated ASVs derived from high-throughput sequencing of cultivated soil bacteria. The inner ring indicates taxonomic placement, while the outer ring shows the soil source of isolation (blue) and denitrification potential (orange) of the corresponding isolates. Denitrification potential was defined as the presence of at least one of the following genes: *narG*, *nirK*, *nirS*, *nosZ* I, or *nosZ* II. The distribution of potential N_2_O reducers is indicated by the asterisk. (b) Number of functional ASVs within the top 9 most diverse genera.

### Keystone taxa associated with denitrification and N_2_O emission in two type of soils

Based on Spearman correlation calculations, microbial co-occurrence networks were constructed for ASVs with more than 50% shared prevalence in both black soil and fluvo-aquic soil samples (r > 0.8, p < 0.05). The prokaryotic microbial communities in black soil and fluvo-aquic soil exhibited different co-occurrence network patterns (Fig. 4a). The co-occurrence network of fluvo-aquic soil was divided into 6 modules, with a modularity value of 0.260, while the black soil co-occurrence network was also divided into 6 modules, but with a higher modularity value of 0.308. Although the number of nodes in both black soil and fluvo-aquic soil networks was similar, the fluvo-aquic soil network had higher average connectivity and average degree (avgK) than the black soil network (Fig. 4b). In the co-occurrence network of black soil, a total of 496 ASVs were identified, among which 2 (0.4%) ASVs served as network hubs. Additionally, 11 (2.2%) ASVs were identified as module hubs, and 173 (34.9%) ASVs were connector hubs (Fig. 4c). In the co-occurrence network of fluvo-aquic soil, a total of 493 ASVs were identified. Among these, 6 (1.2%) ASVs were identified as network hubs via the ZiPi method. Furthermore, 4 (0.8%) ASVs were recognized as module hubs, and 134 (27.2%) ASVs were identified as connector hubs (Fig. 4d).

**Fig. 4.**
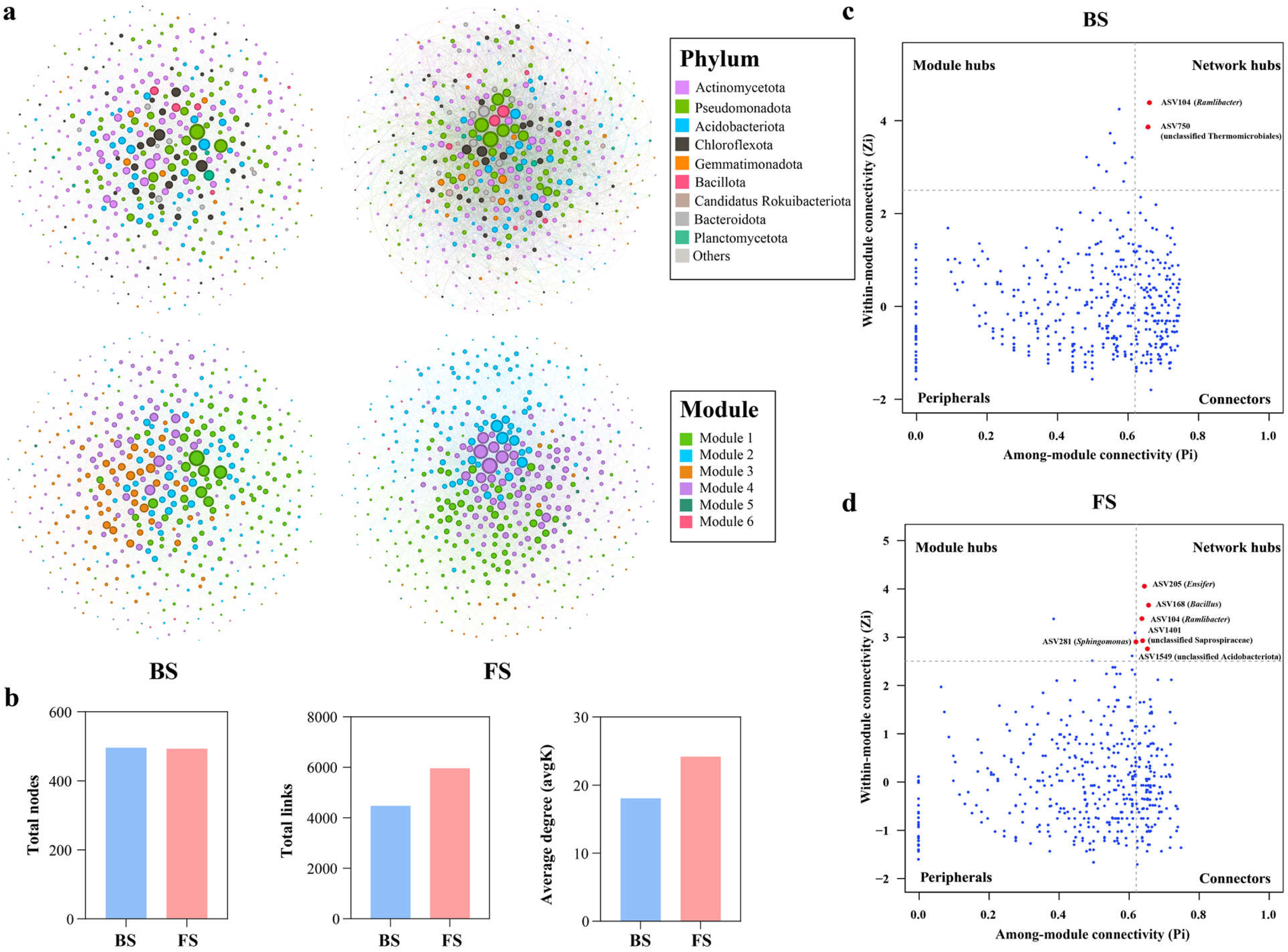
Co-occurrence networks and keystone species of BS and FS soil. (a) Co-occurrence networks of the bacterial community based on pairwise Spearman’s correlations between ASVs. Each shown connection has a correlation coefficient >|0.8| and a P value < 0.05. (b) Total nodes, total links and average degree(avgK) of Co-occurrence networks. (c-d) Zi-Pi plots of keystone taxa in black soil and fluvo-aquic soil based on their topological roles in network. and their community composition at phylum level. Module hubs are identified as Zi ≥ 2.5, Pi < 0.62, connectors are identified as Zi < 2.5, Pi ≥ 0.62, network hubs are identified as Zi ≥ 2.5, Pi ≥ 0.62.

In this study, network hubs characterized by high connectivity both within and between modules were defined as keystone taxa. The potential keystones in black soil included *Ramlibacter* (ASV104) and unclassified *Thermomicrobiales* (ASV750), which belong to the *Pseudomonadota* and *Chloroflexota* phyla, respectively. Compared with black soil, fluvo-aquic soil harbored a significantly higher number of potential key species, primarily belonging to the *Pseudomonadota*, *Acidobacteriota*, *Bacillota*, and *Bacteroidota* phyla. Notable examples included *Ensifer* (ASV205), *Bacillus* (ASV168), *Ramlibacter* (ASV104), *Sphingomonas* (ASV281), unclassified *Saprospiraceae* (ASV1401), and unclassified *Acidobacteriota* (ASV1549). Only *Ramlibacter* (ASV104) was identified as a keystone in both black soil and fluvo-aquic soil. The MAG most closely related with ASV104 is MAG57 *Ramlibacter*, which harbors the majority of the denitrification-associated genes, including *nirK*, *nirS*, *norB*, and *nosZ* gene.

Further statistical evidence from SEM analysis indicates that multiple direct and indirect factors, such as soil properties and nutrients, network hubs, soil community structure, affected soil N₂O emissions. Soil properties and nutrient indicators (pH and DOC/NO₃⁻) and soil community composition directly affect soil N₂O emissions, while the network hubs composition directly impacts the overall soil community structure, subsequently affecting soil N₂O emissions. This ultimately explains 89.6% of the variation in N₂O emissions (Fig. 5a).

**Fig. 5.**
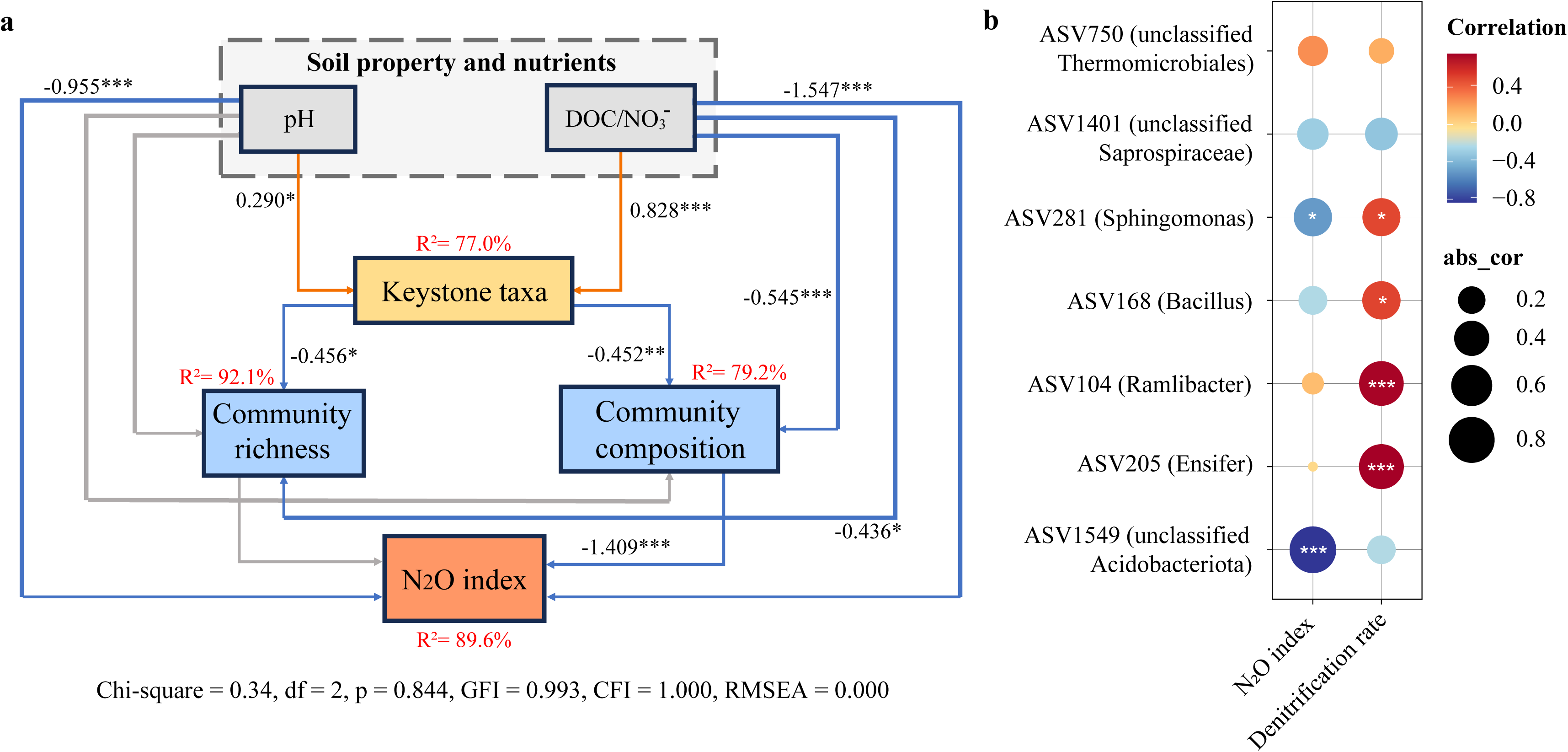
Correlating keystone taxa with potential denitrification rate and N_2_O emission in BF and FS soil. (a) Structural equation modeling of direct and indirect relationships among soil properties and nutrients, network hubs composition, community richness, community composition, and N_2_O emissions. Orange and blue lines represent direct positive and negative effects, respectively; * *p* < 0.05, ** *p* < 0.01, *** *p* < 0.001. The gray line represents no significant; *p* > 0.05. Only significant direct effects are plotted. (b) Correlations between the abundance of keystone species and denitrification rate and N_2_O index of different treatments based on Spearman’s correlation. Significant levels were based on \**p* < 0.05; \*\**p* < 0.01; \*\*\**p* < 0.001.

Spearman correlation calculation was performed based on the relative abundance of key species and denitrification and N₂O emission factors. The relative abundance of *Ramlibacter* (ASV104), *Ensifer* (ASV205), *Bacillus* (ASV168), and *Sphingomonas* (ASV281) showed a significant positive correlation with soil denitrification rates. Among these, *Ramlibacter* (ASV104) (r = 0.74, p < 0.001) and *Ensifer* (ASV205) (r = 0.76, p < 0.001) had the strongest correlations with soil denitrification rates, indicating their significant role in regulating the soil denitrification process. Additionally, the relative abundance of unclassified *Acidobacteriota* (ASV1549) and *Sphingomonas* (ASV281) showed a significant negative correlation with soil N₂O emission factors (Fig. 5b), suggesting their potential involvement in reducing N₂O emissions in soil.

### Comparative genomics analysis of *Ensifer* ASV205 isolates

To gain a deeper understanding of the genomic traits of *Ensifer* ASV205 strains, we selected 10 *Ensifer* strains whose sequences showed 100% identidy to the ASV205 sequence and conducted a comparative analysis of their genomic features. The genome sequencing data of all 10 *Ensifer* ASV205 strains exhibited high quality with the genome sizes ranging from 7.18 to 7.97 Mbp. Each of these strains contain 2 ∼ 5 plasmids. The size of two large plasmids ranges from 1.92 to 2.64 Mbp and 0.67 to 1.67 Mbp. The strains ABS97, AFS42, and OBK33 harbor a third plasmid with varying sizes. ABK11 even has 5 plasmids. Other genomic features of these strains are listed in Supplementary Table S3. When comparing the nucleotide identity (ANI) of these 10 strains, OFS199 showed the greatest difference from the others, with an ANI value ranging from 81.77% to 81.99% (Fig. S3), classifying it as *Ensifer canadensis*. The ANI values among the remaining strains range from 96.6% to 100%, identifying them as *Ensifer adhaerens*.

All 10 *Ensifer* strains possess denitrification functional genes *napA*, *nirK*, and *norB*. Comparative genomics analysis revealed that, except for OFS199, which has only one copy of the *nirK* gene, all other strains contain 2 ∼ 3 copies of the *nirK* gene (Fig. S3a). The *nirK* gene either located on the chromosome and plasmids (Fig. S6). Each strain possesses a single chromosomal copy of the *napA* gene and a single plasmid-borne copy of the *norB* gene (Fig. S6; Fig. S7). Associated with functional phenotypes, strains belonging to group II do not carry the *nosZ* gene, while strains belonging to group Ⅰ and group Ⅲ have one copy of the *nosZ* gene, located on a plasmid (Fig. S3a; Fig. S7). Further comparative genomic analyses revealed pronounced differences in carbohydrate and amino acid metabolism potential among the *Ensifer* ASV205 strains. These genomic variations may contribute to their distinct ecological functions in soil environments (Fig. S4; Fig. S5).

### Divergent **p**henotypic variations in N_2_O metabolism among *Ensifer* ASV205 strains

Through measurements of denitrification and N₂O reduction functions, it was revealed that *Ensifer* ASV205 strains isolated from different soils exhibit distinct capabilities in denitrification and N₂O reduction processes (Fig. 6). Based on the denitrification functional phenotype, we categorized the 10 strains into 3 groups. Group II strains are capable of producing N₂O but lack the ability to reduce it, attributed to the absence of the *nosZ* gene. In contrast, both Group I and Group III strains exhibit dual functions of N₂O production and reduction, though their reduction rates differ significantly. Notably, the difference in denitrification phenotypes between Group I and Group III strains is primarily associated with variations in their growth rates. Although the growth and N₂O reduction rates of OBS65 and OFS103 were similar in DM medium, OFS103 exhibited a higher growth rate compared to OBS65 in TSB medium. After 48 hours of incubation, OBS65 reduced only 1.107 ± 0.370 μmol⋅mL⁻¹ of N₂O-N, while OFS103 reduced 11.25 ± 0.56 μmol⋅mL⁻¹ (Fig. S2).

**Fig. 6.**
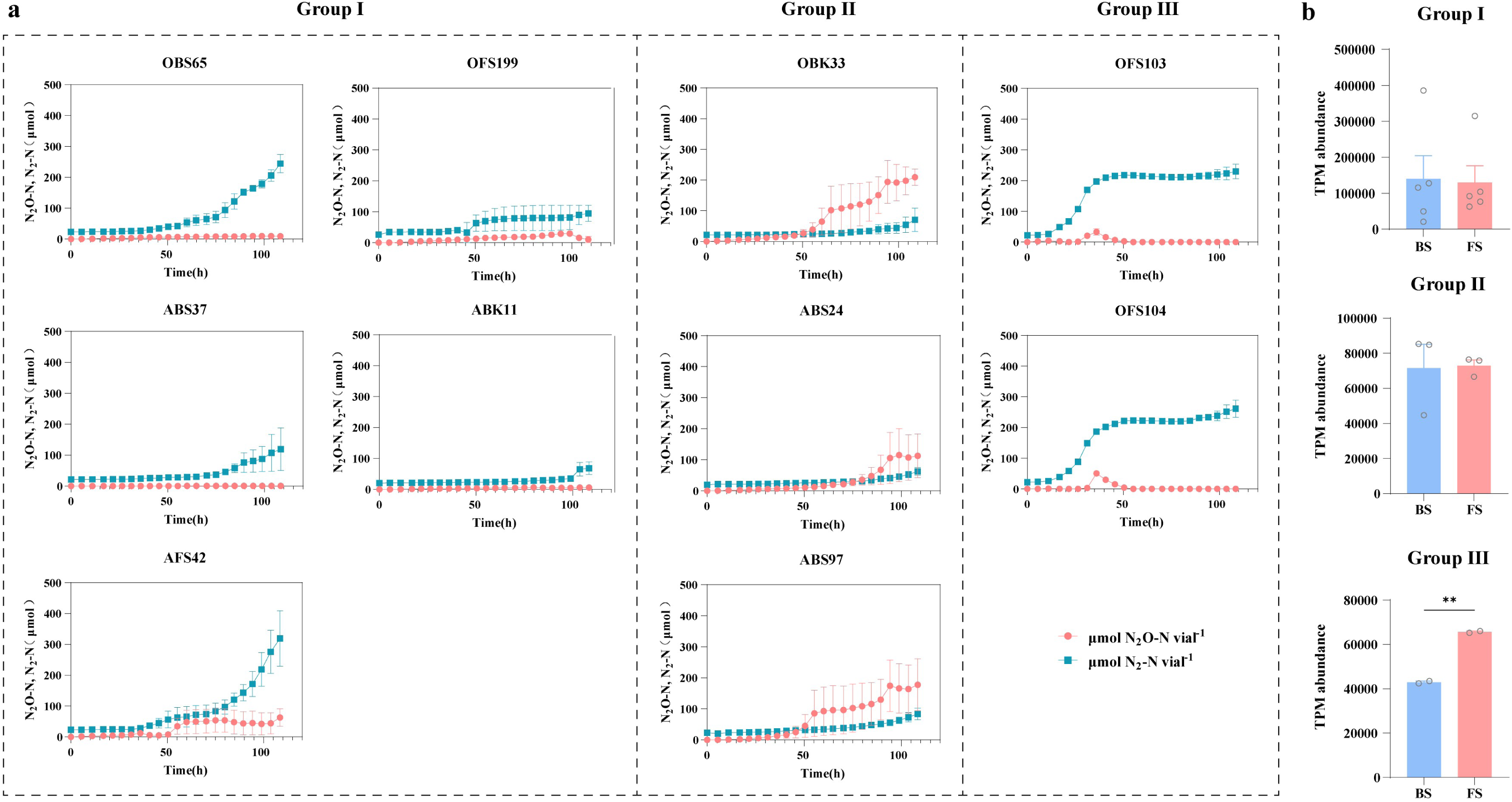
Phenotypic differentiation among three functional groups of *Ensifer* strains and their abundance in two soils. (a) Kinetics of N_2_O and N_2_ in *Ensifer* strains during anaerobic incubation. (b) TPM abundance of *Ensifer* ASV205 strains classified into the three phenotypic groups across BS and FS samples. Paired two-sided t-tests were used. \**p* < 0.05, \*\**p* < 0.01, \*\*\**p* < 0.001.

## DISCUSSION

### Soil-specific patterns in denitrifier diversity and co-occurrence networks

Soil microbial communities play a central role in driving nitrogen transformations and regulating N₂O emissions. Previous studies have shown that the distinct physicochemical properties of black soil and fluvo-aquic soil, shape the composition and function of their respective microbiomes, thereby influencing N₂O dynamics (32, 33). In this study, metagenomic analysis of the species composition in two types of soils revealed that 83.4% of annotated species and 82.6% of annotated denitrifying species were shared between them (Fig. 1b), potentially contributing to overall function of nitrogen cycling. However, significant differences of two soils were observed at the functional level. Although taxonomic composition is largely conserved, FS exhibited significantly higher overall abundance of nitrogen-cycling genes, including those involved in denitrification, nitrification, dissimilatory nitrate reduction to ammonium (DNRA), and assimilatory nitrate reduction. These results suggest that microbial communities in fluvo-aquic soil have higher potential metabolic activity in nitrogen transformations.

However, the net N_2_O emission depends on the balance between microbial N₂O-producing and N₂O-reducing processes. Strain-level variation within shared denitrifying species likely plays a key role in driving soil-specific differences in denitrification capacity and N₂O emissions. It is increasingly apparent that there is no inherent link between species classification and functional potential (34, 35). Within denitrifying microorganisms, even closely related species may exhibit significant differences in functional phenotypes (36). Lycus et al. found that a large number of isolates lacked one or more steps in the denitrification process, and genetic potential cannot always accurately predict actual functional phenotype (31). This difference is mainly attributed to the high diversity of key denitrification functional genes, types of transcriptional regulators, and enzyme assembly (37, 38), which in turn influences their role as either a source or sink of N₂O. Therefore, further exploration of strain-level variation among denitrifiers remains necessary.

Clear soil-specific enrichment of certain taxa was observed at the strain level based on metagenomic and culture-based results. *Castellaniella* MAGs carrying *nirK* and *nosZ* genes were enriched in FS, and their isolates were obtained exclusively from FS samples (Fig. 2; Fig. 3). *Castellaniella* has been reported to play a role in reducing soil N₂O emissions (32). *Ensifer, Mesorhizobium* and *Cupriavidus* showed high strain-level diversity, with the majority of strains capable of denitrification and N_2_O reduction. The isolates of *Ensifer* and *Cupriavidus* were obtained from both soils in this study, while the majority of *Mesorhizobium* isolates primarily isolated from black soil. These potential denitrifying bacteria and their differentiated distribution may contribute to functional variation of denitrification in two different soils.

Beyond compositional differences, ASV-based co-occurrence network analysis further revealed distinct organizational strategies among these microbial taxa. The fluvo-aquic soil network exhibited substantially higher average degree and connectivity, reflecting tighter microbial associations. This denser network may facilitate more rapid resource exchange and collective responses to environmental fluctuations, potentially contributing to the higher abundance of functional gene observed in FS (39, 40). Compared to fluvo-aquic soil, black soil exhibits a higher network modularity value, indicating a more structured organization within the soil microbial community. These modules comprise members with similar functions, which can carry out specific tasks such as the nutrient cycling of carbon, nitrogen, and other elements (41). This may reflect adaptation to the spatially heterogeneous but organic-rich environment of black soil, which promotes localized nutrient cycling strategies (42).

### Potential contribution of keystone taxa to denitrification and N₂O emission

In this study, key ASVs with high connectivity in the microbial co-occurrence network were identified as keystone taxa, although they are not the top dominant ASVs within the microbial community. Studies have shown that keystone species not only influence other species and regulate overall community structure through the secretion of metabolites, antibiotics, or toxins (43), but also act as functional species that enhance the availability of resources essential for nutrient cycling and facilitate the development of microbial networks (7, 44). In this study, SEM provided evidence that these keystone ASVs, together with environmental factors, collectively explained most of the variation in soil N₂O emissions. These keystone ASVs likely influence N₂O emissions by structuring the microbial community. However, their ecological impact appears to be context-dependent, as their activity and network centrality are shaped by environmental conditions such as pH and carbon/nitrogen availability.

Among the identified ASVs, *Ramlibacter* ASV104 was identified as a keystone species in both type of soils (Fig. 4; Table S2). This ASV showed close correlation with soil denitrification rates. Metagenomic assembly results confirmed that the closest MAG of *Ramlibacter* ASV104 possesses denitrification potential for N₂O production and reduction. Previous studies have reported that *Ramlibacter* exists in a wide range of ecosystems, including deserts (45), soils (46), and freshwater (47), indicating that it possesses an ecological advantage across a broad range of ecological niches. Studies focusing on marine and soil environments have revealed that *Ramlibacter* is closely linked to the denitrification process and N₂O emissions (33, 48).

In FS, *Ensifer* ASV205 exhibited high node degree, betweenness centrality, and node stress, positioning it as a keystone ASV in the microbial network (Table S1). Ten isolates matching the 16S rRNA sequence of ASV205 were obtained from both BS and FS, and all exhibited denitrification capabilities. Interestingly, these strains displayed notable phenotypic divergence under different cultivation conditions, indicating their potential to display adaptive and divergent functional traits in the heterogeneous soil environment. Most of *Ensifer* isolates in this study belonged to *Ensifer adhaerens*, which is commonly found in soil and rhizosphere environments for their ability to promote plant growth, fix nitrogen (49), and denitrification abilities (50, 51). Notably, previous studies have shown that keystone taxa in soil often possess a high copy number of functional genes related to carbon, nitrogen, phosphorus, and sulfur cycling (12). Genomic analyses of *Ensifer* ASV205 strains revealed multiple copies of the *nirK* gene distributed across chromosomes and plasmids (Fig S6), potentially enhancing their nitrite tolerance and adaptation to variable environmental conditions. One study have found that *Ensifer* strains with different copies of functionally redundant genes and varied locations of these genes within the genome exhibit differential sensitivity in response to changing environmental conditions (52). Such variation in gene copy number and genomic location, together with differences in the ability to utilize carbon sources, amino acids, and other nutrients, may drive the ecological differentiation of *Ensifer* ASV205 strains within varied soil environments, ultimately shaping their distinct roles in denitrification processes and N₂O emissions.

Metagenomic read mapping indicated that a group of *Ensifer* ASV205 (Group III) were more abundant in fluvo-aquic soil, likely due to their enhanced growth potential in this type of soil due to different nutrient utilization strategy. In soils with fluctuating nutrient availability, bacterial growth rate plays a key role in adaptation and resource allocation (53). These traits may enable certain *Ensifer* strains to establish dense metabolic interactions with other community members in fluvo-aquic soil, thereby facilitating their emergence as network hubs. This hub species in the microbial community may further play a key role in shaping the whole community structure as demonstrated in previous literature (54). In contrast, within the distinct environmental context of black soil, *Ensifer* ASV205 lost its centrality, with other denitrifiers like *Ramlibacter* emerging as key keystone taxa in the microbial network.

## MATERIALS AND METHODS

### Soil sampling

Black soil (BS) was collected from long-term experimental fields in the Northeast Plain and fluvo-aquic soil (FS) was obtained from the North China Plain. Samples were collected using five-point sampling method at depths of 0-20 cm and homogenized to form composite samples. Soil samples were sieved through a 2.0-mm mesh to remove debris and stored at 4°C until further use. The detailed physicochemical properties of the soil were presented in previous study (32). To stimulate denitrifying bacterial activity, 20 g (dry weight) of each soil sample was placed into 120 mL serum bottles and adjusted to 70% water-holding capacity. The bottles were sealed with butyl rubber stoppers and aluminum caps, and the headspace was flushed with high-purity helium (99.999%) by alternately evacuating and refilling four times to establish an anaerobic environment. Nitrate (250 mg/kg) and glucose (1,000 mg/kg) were added to stimulate the denitrifying microorganisms. All bottles were incubated at 25 °C for 7 days in triplicates. The enriched soil samples were subsequently used for both metagenomic sequencing and bacterial isolation.

### Metagenome sequencing and data analysis

Soil samples were collected for DNA extraction with the Omega Bio-Tek soil DNA kit and library construction using the Illumina TruSeq DNA sample preparation guide. Each library was sequenced by Illumina NovaSeq platform (Illumina, USA) with PE150 strategy at Personal Biotechnology Co., Ltd. (Shanghai, China). Raw sequencing reads were processed to obtain quality-filtered reads for further analysis. Specifically, fastp (v0.23.2) (55) was employed to perform quality control and adapter trimming. After the quality control process, we obtained 98.6 gigabases (Gb) of metagenomic clean reads, with average value of 16.4 Gb data per sample (range from 15.0 to 17.0 Gb). Filtered reads were then assembled to contigs using megahit (v1.1.2) (56). Prodigal (v2.6.3) (57) was used to predict the genes in the contigs. To assess the abundances of these genes, the high-quality reads from each sample were mapped onto the predicted gene sequences using Minimap2 (58) and using featureCounts to count the number of reads aligned to gene sequences, and normalized to transcript per million (TPM) based on the gene length and sequencing depth. Nitrogen cycle-related functional genes were annotated using Diamond (v2.0.15) against NCycDB database.

Metagenome assembled genomes (MAG) were constructed from the shotgun metagenome data. Assembled contigs were further binned to metagenome-assembled genomes using metaBAT2 v2.12.1(59). The quality of the MAGs were assessed via the lineage-specific workflow of CheckM v1.2.3(60), and those with <50% completeness or >10% contamination were excluded. The pairs of MAGs with >99% ANI were dereplicated using the ‘dereplicate’ command in dRep v3.5.0 (61). The TPM abundance of MAGs was calculated using CoverM (v0.7.0) (62). Taxonomy annotation of the MAGs was performed using the GTDB-Tk (v 1.0.2). The phylogenetic tree based on 120 bacterial universal marker gene set generated by GTDB-Tk was visualized using the Interactive Tree of Life (iTOL) online tool (https://itol.embl.de/). Nitrogen cycle-related functional genes were annotated using Diamond (v2.0.15) against NCycDB database.

### Microbial co-occurrence network construction and keystone taxa identification

We collected 16S rRNA sequencing data of 9 black soil and 9 fluvo-aquic soil samples from previous research (32). All steps of sequence processing and quality control were performed in QIIME2 (v2021.4) [35]. Representative sequences for each ASVs were assigned taxonomic classifications using the SILVA132 16S rRNA database [36]. To ensure comparability, 16S rRNA sequences were rarefied to the 11000 sequencing depth. The construction of microbial co-occurrence networks for each soil type was constructed by using the Integrated Network Analysis Pipeline (iNAP, http://mem.rcees.ac.cn:8081) (63). Robust correlations, with Spearman correlation coefficients (ρ) >0.8 or <−0.8 and FDR-adjusted p-values <0.05, were used to construct the networks. Network complexity was estimated by calculating features such as the total number of nodes, links, average degree and modularity. The networks were visualized using Gephi v0.10.1 software.

To identify potential keystone species in the networks of black soil and fluvo-aquic soil, the within-module connectivity (Zi) and between-module connectivity (Pi) of each node were calculated. Nodes with high values of Zi or Pi (i.e., keystone taxa) were defined as module hubs (Zi > 2.5, Pi ≤ 0.62), network hubs (Zi > 2.5, Pi > 0.62), and connectors (Zi ≤ 2.5, Pi > 0.62) (64, 65).

### Isolation, high-throughput sequencing, and functional identification of isolates

Dilute three grams of black soil and fluvo-aquic soil in 0.85% saline at a ratio of 1:10. The diluted soil suspension was then serially diluted and spread individually onto each of the five replicate plates with 1/10 TSA medium (Merck, Germany). The plates were incubated in parallel under both aerobic and anaerobic conditions at 28°C. All individual colonies with distinct morphologies were selected and purified to obtain the most diverse bacterial isolates possible. A total of 719 isolates were obtained. The lysis buffer containing 1% Triton X-100 and 0.5% Tween-20 was used for rapid DNA extraction from bacterial isolates. The primers targeting the *narG*, *nirK*, *nirS*, *nosZ* Ⅰ and *nosZ* Ⅱ gene were used for screening the potential denitrifiers and N_2_O reducers. Primer sequences and corresponding information were summarized in Supplementary Table S4. The taxonomy of all isolates was determined using a high-throughput sequencing approach, as previously described (66). Detailed protocols for high-throughput identification and phylogenetic analysis of isolates are provided in Supplementary Materials and Method.

To compare the denitrification capabilities of different *Ensifer* ASV205 strains, 30 mL of tryptic soy broth (TSB) medium was inoculated with bacterial isolates and incubated under anaerobic conditions. Sodium nitrate (NaNO₃, 5.9 mM) and sodium nitrite (NaNO₂, 1.5 mM) were supplemented as terminal electron acceptors in the medium. To compare the growth rate and N₂O reduction rate of *Ensifer* ASV205 strains OBS65 and OFS103, the strains were cultured in 30 mL of TSB medium and nitrogen-free DM medium. N₂O (204.5 μmol) was injected into the headspace of the serum bottles using the sterile syringe. The N_2_O and N_2_ concentration in the headspace was measured using the robotized incubation system (67).

### Whole genome sequencing and assembly

The strains were cultured overnight in TSB medium. The genomic DNA was extracted by using the Cetyltrimethyl Ammonium Bromide (CTAB) method with minor modification, and then the DNA concentration, quality and integrity were determined by using a Qubit Flurometer (Invitrogen, USA) and a NanoDrop Spectrophotometer (Thermo Scientific, USA). Sequencing libraries were generated using the TruSeq DNA Sample Preparation Kit (Illumina, USA) and the Template Prep Kit (Pacific Biosciences, USA). The genome sequencing was then performed by Personal Biotechnology Company (Shanghai, China) by using the Nanopore PromrthION48 platform and the Illumina Novaseq platform. Data assembly was proceeding after adapter contamination removing and data filtering by using AdapterRemoval (68) and SOAPec (69). The filtered reads were assembled by SPAdes (70) and A5-miseq (71) to constructed scaffolds and contigs. The average nucleotide identity (ANI) index was calculated using JSpecies v1.2.1 software (72). Gene prediction was performed by GeneMarkS v4.32 (73). CAZy (Carbohydrate-Active enzymes) database (74) was used to predict Carbohydrate-Active enzymes. Function annotation was completed by blast search against different KEGG (Kyoto Encycolpedia of Gene and Genomes) database (75). MEGA11 (76) software was used to construct the phylogenetic tree. To estimate the in-situ abundance of genomes, TPM tables were generated using coverm v0.7.0 (62). Phylogenetic analysis of denitrification genes of the *Ensifer* ASV205 isolates is provided in Supplementary Materials and Method.

### Bioinformatic and statistical analyses

Based on the 18 soil samples and their corresponding physicochemical properties determined in the previous study (32), structural equation modeling (SEM) was conducted using IBM SPSS Amos 26 (IBM Corporation, NY, USA) to investigate the hypothesized relationships between microbial community structure and N₂O emission patterns (65, 77). R v4.1.3 was used for performing the spearman correlation calculation and visualizing the results.

## Data availability

All nucleotide sequences obtained in this study have been deposited in GenBank. Amplicon sequences from the 16S rRNA gene survey were deposited in NCBI BioProject PRJNA755188. Metagenomes, MAGs and the complete genome sequences of *Ensifer* strains are available under BioProject PRJNA1283983.

## ACKNOWLEDGMENTS

This work was supported by the National Natural Science Foundation of China (NSFC 31971526), and the Key R&D project of Ministry of Science and Technology (2017YFD0200102).

## AUTHOR CONTRIBUTIONS

Siyu Yu: Investigation, Software, Methodology, Data curation, Formal analysis, Visualization, Writing – original draft, Writing – review & editing. Qiaoyu Wu: Investigation, Methodology, Data curation. Yiming Ma: Methodology, Data curation. Xiaojun Zhang: Resources, Conceptualization, Writing – review & editing, Supervision, Funding acquisition.

## CONFLICTS OF INTEREST

The authors declare that they have no competing interests.

## SUPPLEMENTAL MATERIAL

Supplemental methods, results, tables and figures. PDF file, 1.29 MB.

